# Comprehensive in situ mapping of human cortical transcriptomic cell types

**DOI:** 10.1101/2021.01.28.428548

**Authors:** Christoffer Mattsson Langseth, Daniel Gyllborg, Jeremy A. Miller, Jennie L. Close, Brian Long, Ed S. Lein, Markus M. Hilscher, Mats Nilsson

## Abstract

The ability to spatially resolve the cellular architecture of human cortical cell types over informative areas is essential to understanding brain function. We combined in situ sequencing gene expression data and single-nucleus RNA-sequencing cell type definitions to spatially map cells in sections of the human cortex via probabilistic cell typing. We mapped and classified a total of 59,816 cells into all 75 previously defined subtypes to create a first spatial atlas of human cortical cells in their native position, their abundances and genetic signatures.

## INTRODUCTION

The human cortex contains roughly 80 billion cells^(1,2)^ and understanding the identity of all the cells is challenging. Evolving single-cell technologies have recently allowed scientists to start to comprehend the cellular composition of the human cortex^(3–5)^, enabling quantitative descriptions of cellular diversity and definitions^(6)^. Hodge and colleagues characterized 15,928 cell nuclei from human middle temporal gyrus (MTG, Brodmann area 21, a part of the temporal lobe) by single-nucleus RNA-sequencing (snRNA-seq) ^(4)^. However, the precise laminar locations and abundances of many of these cell types have not yet been described, and beyond coarse layers most of these types are at low proportions and intermingled with one another. Hybridization-based *In Situ* Sequencing (HybISS) is an image-based multi-targeted gene expression profiling technique that allows the precise mapping of individual cells in human brain tissues^(7)^. Various analytical approaches can assign detected transcripts to segmented cells and subsequently, cells to cell types. One such approach, probabilistic cell typing by *in situ* sequencing (pciSeq), leverages single-cell RNA-sequencing data to guide cell type assignment^(8,9)^. Here, we implement pciSeq to map cell types across three human cortical sections as a proof of principle to show an efficient and robust method to accurately resolve anatomical organization of human tissue that is envisioned for such efforts as the Human Cell Atlas^(10)^.

## RESULTS

As part of a Human Cell Atlas pilot project to explore cell type mapping with spatial transcriptomics methods (the SpaceTx consortium), we obtained human temporal lobe tissue from surgical resections (**Figure 1a**). Using a panel of 120 genes chosen to span snRNA-seq cell type definitions in the middle temporal gyrus, we applied pciSeq to produce cellular maps of human brain tissue. We mapped 59,816 cells of 75 transcriptomic cell types including 24 glutamatergic, 45 GABAergic and 6 non-neuronal cell types (Section A: 19,127 cells, 27.51 mm^2^; Section B: 28,694 cells, 41.41 mm^2^; Section C: 11,995 cells, 30.23 mm^2^; **Figure 1a, b; Supplementary Figure 1**). pciSeq assigns cells, represented as pie charts, with the angle of each slice proportional to the cell type probability. For instance, a cell of subclass Layer 2/3 can have 72.8% probability of being an Exc L2-3 *LINC00507 FREM3* cell type and 27.2% being an Exc L2-3 *GLP2R* cell type (**Figure 1c; Supplementary Figure 2**). Here, the highest probability in the pie chart defines the final cell type for all downstream analysis. The level of certainty for each cell subclass is shown in **Supplementary Figure 3**. The size of each pie chart is indicative of the number of assigned transcripts. On average, 70.9 ± 7.4 transcripts and 30.4 ± 2.0 distinctive genes were measured in neuronal cells, approximately twice as many as non-neuronal cells (**Supplementary Figure 4**). Counting the cell type occurrences in our tissue sections shows that non-neuronal cell types outnumber neuronal cell types by 3.38, comparable to published results that measured a ratio of 3.76 over the entire human cortex^(1)^.

**Figure 1:**
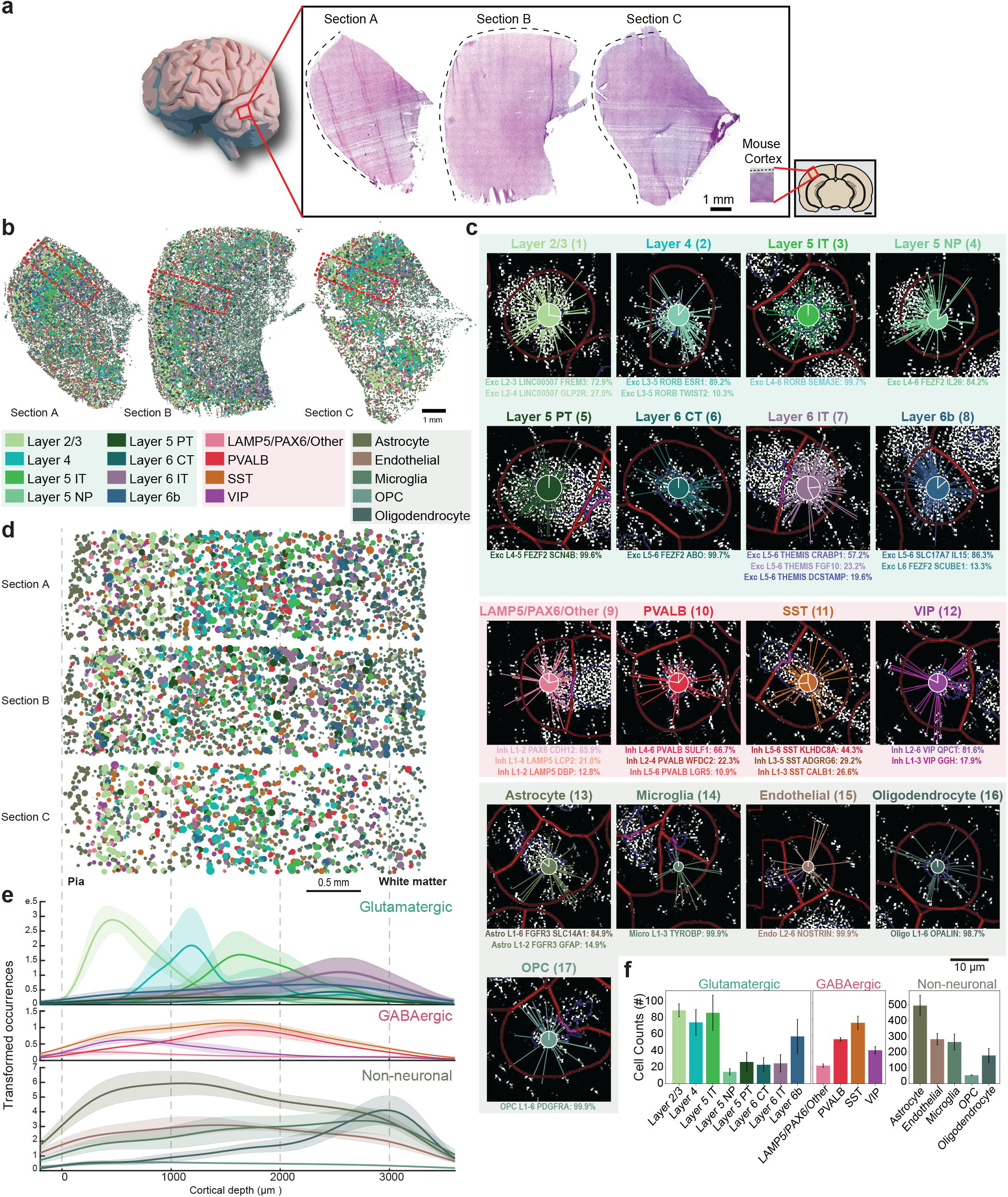
Cell types of the human temporal lobe. (**a**) Three tissue sections from the temporal lobe (Sections A-C) and the mouse visual cortex for size reference. (**b**) Spatial maps of cell types across human temporal lobe (Sections A-C), including 24 glutamatergic, 45 GABAergic and 6 non-neuronal cells. The cells are represented as pie charts where the angle of each slice is proportional to the likelihood of the cell being of a certain type and the size of the pie chart is indicative of the number of transcripts. The colors correspond to the cell subclasses stated below the maps. Red rectangles mark the regions of interest (ROIs) in (d). (**c**) Genes are assigned to cells and cells are subsequently classified based on single-nucleus RNA-sequencing (snRNA-seq) data. Examples shown are median cells for each cell subclass, i.e. cells with the median count of the number of transcripts being assigned within each subclass. The number next to the cell subclass label denotes the cell location in Supplementary Figure 2. The probability for each cell type is listed below the example cells. (**d**) ROIs spanning across the neocortical layers in the tissue sections in (a). (**e**) Mean cortical depth profiles from the ROIs, with the transparent shades representing the standard error of the mean. Y-axis represents occurrences after smoothening data with a kernel (100 bins). Separated histograms for each cell subclass are shown in Supplementary Figure 6. (**f**) Mean counts of each cell subclass found within the ROIs (error bars represent standard error of the mean).

To accommodate for tissue variability, regions of interest (ROIs) were outlined spanning the six neocortical layers and some white matter (**Figure 1d; Supplementary Figure 5a**). First, we investigated the distribution of the assigned cell types as a function of cortical depth, measured from the pial surface (**Figure 1e; Supplementary Figure 6**). Glutamatergic cells often showed characteristic depth profiles corresponding to cortical layers while GABAergic cells were less confined. However, the *VIP* subclass and the *LAMP5*/*PAX6*/Other subclass had the highest density in supragranular layers, whereas the *PVALB* subclass and *SST* subclass peaked around the granular layer, similar to what has been shown in mouse neocortex^(11,12)^. Among the non-neuronal cells, oligodendrocytes were most distinct, showing a higher cellular density in infragranular layers and white matter. Counting the occurrence of neuronal cell types showed a proportion of 67:33 for glutamatergic versus GABAergic cells. Layer 2/3, Layer 4, Layer 5 IT and Layer 6b cells were the most abundant cell types in the glutamatergic population and *SST* and *PVALB* cells in the GABAergic population (**Figure 1f**).

To further examine the neocortical architecture, we manually demarcated the six cortical layers guided by known layer markers (*LAMP5*, *LINC00507*, *COL5A2*, *FEZF2*) and estimation of published layer thickness^(13)^ (**Figure 2a,b; Supplementary Figure 5b, 7 and 8a**). We quantified the ratio of cell types within and across layers. Within-layer distribution of glutamatergic cells largely followed the proposed layer locations of Hodge *et al*.^(4)^ (**Figure 2c**). For example, 79% of mapped cells in layer 2 were classified as Layer 2/3 cells whereas 53.7% of mapped cells in layer 4 were classified as Layer 4 cells. Compared to glutamatergic neurons, GABAergic neurons (**Figure 2d**) and non-neuronal cells (**Supplementary Figure 8b**) were more homogeneously distributed. These trends followed the snRNA-seq data (**Supplementary Figure 9a,b,c**). Across layers, we further separated the eight glutamatergic subclasses into the 24 cell types, the four GABAergic subclasses into 45 types and the five non-neuronal subclasses into 6 types (**Figure 2e,f; Supplementary Figure 8c**). Glutamatergic cell types of supragranular and granular layers (L1-3 and L4 respectively) showed clear layer structure (**Figure 2e**), and similar across-layer distributions as the snRNA-seq data (quantified with Pearson correlation coefficient; **Supplementary Figure 9f and 10a**). The mean correlation coefficient was 0.77 ± 0.05 for glutamatergic cells. Considering the layer-specificity of the genes that define these cell types, we noted that cells in supragranular and granular layers were well defined by the genes *LINC00507* and *COL5A2* respectively and infragranular layers (L5-6) were mainly defined by *FEZF2* (**Figure 2g; Supplementary Figure 11a**). GABAergic cell types were sparser than glutamatergic cell types, with *LAMP5*/*PAX6*/Other and *VIP* cells escalating in supragranular layers and *SST* and *PVALB* cells around the granular layer (**Figure 2f**). Comparing the differences in layer distribution of cells mapped with pciSeq and snRNA-seq, the mean Pearson correlation coefficient was 0.63 ± 0.04 (**Supplementary Figure 9e and 10b**). *LAMP5* cell types, in particular, followed the proposed cell type location by Hodge *et al*.^(4)^ Also, *ADARB2* (a marker for CGE-derived cells, such as *LAMP5* and *VIP* cells) and *LHX6* (a marker for MGE-derived cells, such as *PVALB* and *SST* cells) showed a clear separation between cells in supragranular layers versus cells in infragranular layers (**Figure 2h**). Similar to glutamatergic cells, marker gene expression was not entirely restricted to layers. While *VIP* gene expression was most abundant in supragranular layers, it could also be found in infragranular layers (**Supplementary Figure 11b**). Among the cell types in the non-neuronal population, there was a notable increase in the density of cells in infragranular layers and white matter (**Supplementary Figure 7c and 8a**). The oligodendrocytes (Oligo L1-6 *OPALIN*) exhibited a distinct layer distribution being more abundant in the infragranular layers (**Supplementary Figure 8c**). This was consistent with the principal marker for the oligodendrocytes, *OPALIN* as it was mostly expressed in the infragranular layers and white matter (**Supplementary Figure 8d and 11c**). Quantifying the difference in layer distribution between pciSeq and snRNA-seq showed lower agreement (0.35 ± 0.18 mean Pearson correlation coefficient) to the snRNA-seq (**Supplementary Figure 9f and 10c**).

**Figure 2:**
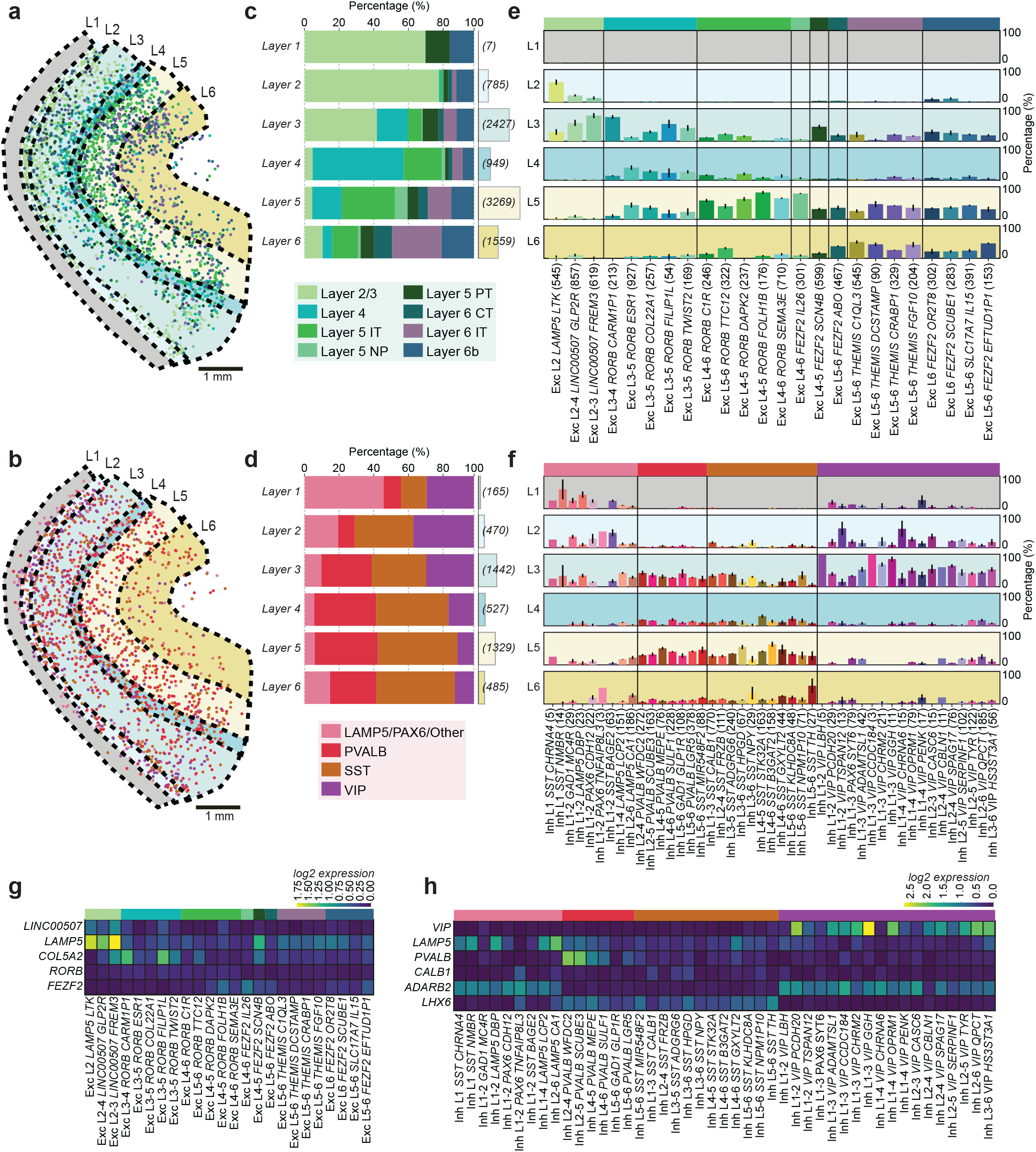
Layer-specificity and molecular composition of neurons in the human temporal lobe. (**a**) Location of glutamatergic cells colored by the most probable cell subclass and annotated neocortical layers (L1-6). (**b**) Same as in (a) for GABAergic cells. (**c**) The within-layer relative distribution of the glutamatergic cells and the number of cells counted for each layer in brackets. (**d**) Same as (b) for the GABAergic cells. (**e**) Across-layer distribution of glutamatergic cell types. The colored bars represent the relative proportion of each cell type in each layer (L1-6). (**f**) Same as (e) for GABAergic cell types. (**g**) Mean log2-transformed expression of known glutamatergic marker genes. (**h**) Same as (g) for GABAergic marker genes.

## DISCUSSION

Taken together, we mapped 24 glutamatergic, 45 GABAergic and 6 non-neuronal cell types previously classified by snRNA-seq^(4)^. Within the targeted gene list, classical markers for glutamatergic, GABAergic and non-neuronal cell types were included to aid in probabilistic assignment of cells *in situ*. However, some classical markers such as *SST*, a main marker in the human cortex, was not prioritized by the gene selection approach. Therefore, *SST* subclass genes were used for cell calling. The combinatorial detection approach of the genes allows and assures the precise assignment of molecularly defined cell classes as well as their subtypes *in situ*.

Our spatially resolved, transcriptomically profiled tissue sections display a similar non-neuronal to neuronal ratio as the previously reported, immunocytochemically measured ratio in the entire cerebral cortex^(1)^. Comparison to published sequencing data also suggests high correlation in the layer distribution between *in situ* data and snRNA-seq^(4)^ data. Here, manual layer annotations were guided by four known marker genes but layer outlines could also be identified by cellular granularity or more automated approaches^(14,15)^ which take into consideration that layer boundaries are not sharp. We found mainly glutamatergic cell types to arrange in layers but they were less layer-restricted than expected^(4)^, advocating that anatomical location alone is not sufficient to characterize a cell.

pciSeq presents itself as a powerful tool in cell type assignment across large tissue areas, not only for *in situ* sequencing data but also other spot-based spatial approaches^(16,17)^. The presented human cortical maps include a comprehensive reference of the cells in the human temporal lobe, their spatial location, abundances and gene expression signatures, all central features outlined by the Human Cell Atlas^(10)^ and others^(6)^. This work embodies the vision and paves the path towards a spatial Human Cell Atlas^(10)^, utilizing the predefined taxonomy of cells^(4)^ to create maps of histological tissue structures.

## METHODS

### Tissue preparation

Surgically resected temporal lobe tissue was divided into 500 μm slices using an EMS TC-1 tissue chopper. The 500 μm sections were then further sliced to 10 μm and collected on slides. Slices were placed on the flat bottom surface of a cryo embedding mold and embedded in OCT. After 15 minutes equilibration in OCT, slices were frozen in a dry ice - ethanol slurry, vacuum sealed and stored at −80°C.

### Tissue description

Section A and C are from the middle temporal gyrus of a 38-year old epilepsy donor. Section B is from the fronto-temporal parietal lobe of a 72-year old male tumor case.

### HybISS

Technical details and description of the HybISS method can be found published in Gyllborg *et al*.^(7)^ as well as transcript identity and coordinates for Sections A and B. The same gene panel and experimental procedures were used in the generation of Section C. The genes targeted in the HybISS experiments were manually and computationally curated as a part of the Chan Zuckerberg Initiative SpaceTx Consortium. The panels and the computational selection were based on snRNA-seq data from human middle temporal gyrus^(4,18)^.

### Image analysis

The description of the image analysis can be found in Gyllborg *et al*.^(7)^ The same in-house software was used for the generation of the spatial gene expression data and is available at <https://github.com/Moldia>.

### Cellular segmentation

Cells were segmented using CellProfiler 2.2.0^(19)^ in which the diameter of the objects were set to fit the span of nuclei sizes. The DAPI images were tiled to reduce the computational requirements. A manual threshold was set for localizing the nuclei. The objects localized were then expanded and subsequently converted to images. The area and location of the objects were recorded.

### pciSeq

The pciSeq pipeline can be found at <https://github.com/acycliq/full_coronal_section> and is described in Qian *et al*.^(8)^. In short, the pciSeq pipeline works by assigning genes to cells and then cells to cell types. This assignment is done using a probabilistic framework based on single-cell RNA sequencing data. The input data required to run the algorithm consists of the area, the global x and y coordinates and the unique cellular identifier of the segmented cells. In addition, global x and y coordinates of the transcript, the transcript type and the cell that the transcript belongs to is needed for each transcript. Cell type definitions used in pciSeq were downloaded from the Allen Brain Atlas <https://portal.brain-map.org/atlases-and-data/rnaseq>. All of the colors used in the figures follow the cell type color definition formulated by the Allen Institute.

### pciSeq processing module

The downstream processing of the data was done using a set of functions created in-house <https://github.com/Moldia/pciseq_processing_module>. The set of tools included finding the most probable cell type call from the pciSeq output, plotting of these cell type calls, plotting the cell types individually, adding polygon labels to the cells and creating cell-by-gene matrices of the called cells.

### Region of interest and layer annotations

The regions of interest (ROIs) were drawn using napari^(20)^, a multi-dimensional image viewer for python. The ROIs were outlined to cover the six neocortical layers and some white matter. The area of each ROI was the same for all tissue sections (3.2 mm^2^). The layers were annotated in napari based on the expression of known marker genes (*LAMP5*, *LINC00507*, *COL5A2*, *FEZF2*) in addition to the relative thickness of cortical layers^(13)^.

### Software

The code used to generate the plots for the Figures is available at <https://github.com/Moldia/spatial_mapping_of_human_cortical_cells>.

## ACKNOWLEDGMENTS

We thank all members of the SpaceTx Consortium. This publication is part of the Human Cell Atlas <https://www.humancellatlas.org/publications>. We thank our funding agencies for making this work possible: Chan Zuckerberg Initiative, an advised fund of Silicon Valley Community Foundation; Erling-Persson Family Foundation; Knut and Alice Wallenberg Foundation; Swedish Research Council [2019-01238]; Swedish Brain Foundation (Hjärnfonden) [PS2018-0012].

## Supplementary Materials

Supplementary materials contain 11 figures.

