## Supplementary Materials for "Comprehensive in situ mapping of human cortical transcriptomic cell types"

Supplementary materials contain 11 figures.

### Supplementary Figure 1

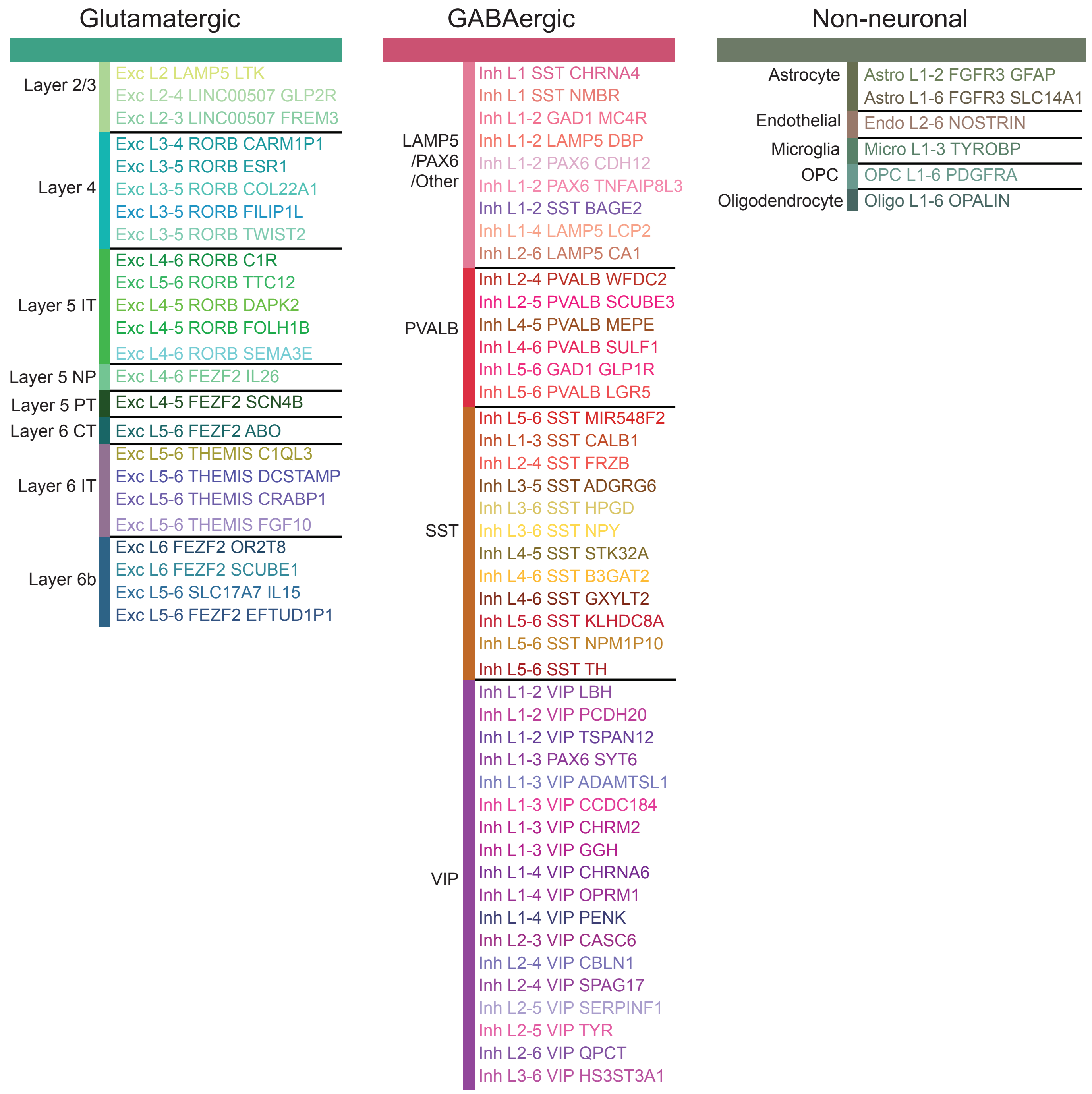


**Supplementary Figure 1 | Cellular hierarchy and cell type definitions.**

The top level is the separation into glutamatergic, GABAergic and non-neuronal cells. The second level is the separation of the glutamatergic, GABAergic and non-neuronal cells into 17 subclasses. The third level is the separation of the 17 subclasses into 75 cell types.

#### Supplementary Figure 2

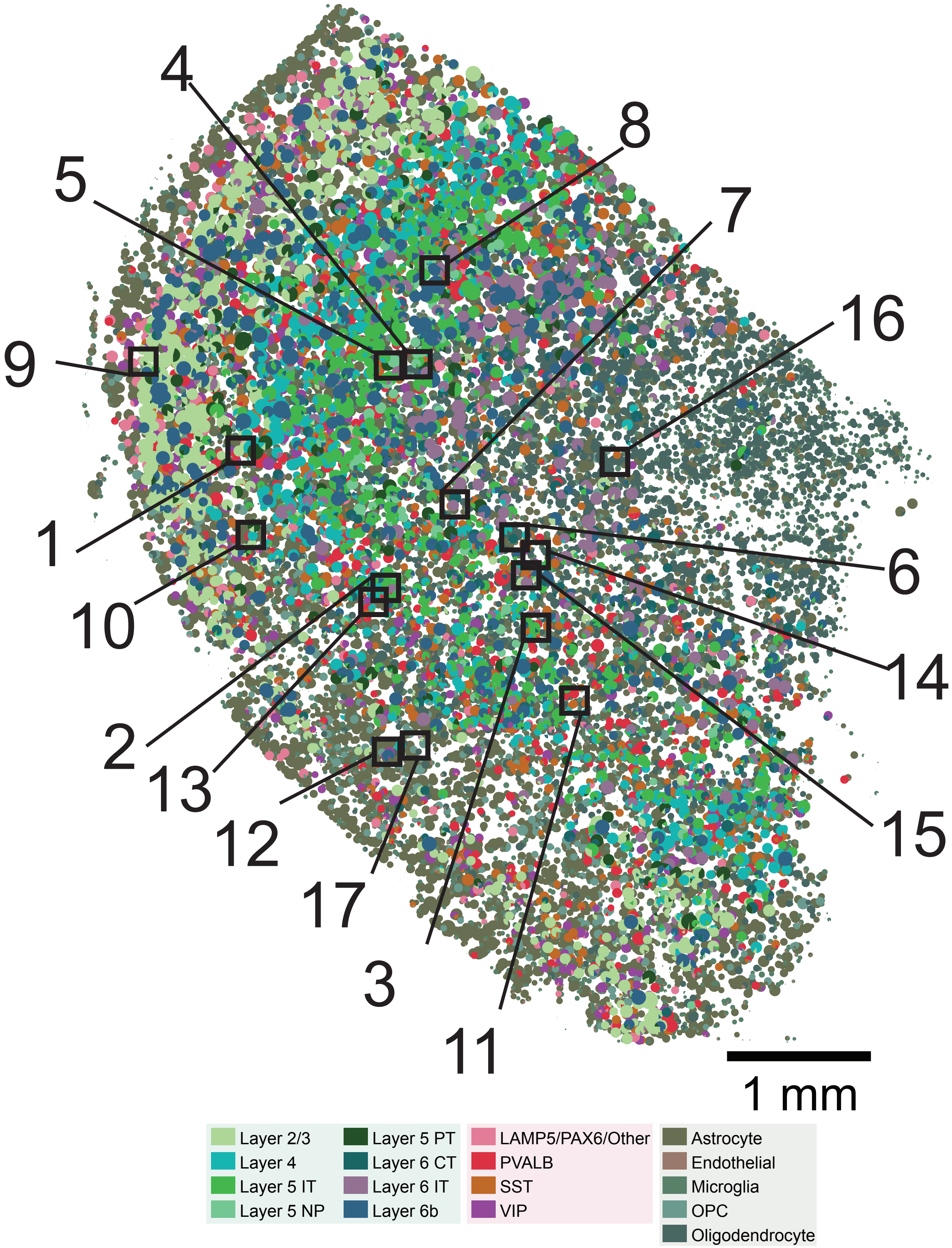


**Supplementary Figure 2 | Approximate location of example cortical cells.**

The location of the example cells shown in Figure 1b. The colors correspond to the 17 cell subclasses. Black boxes represent the approximate location of each cell and the number denotes the corresponding cell.

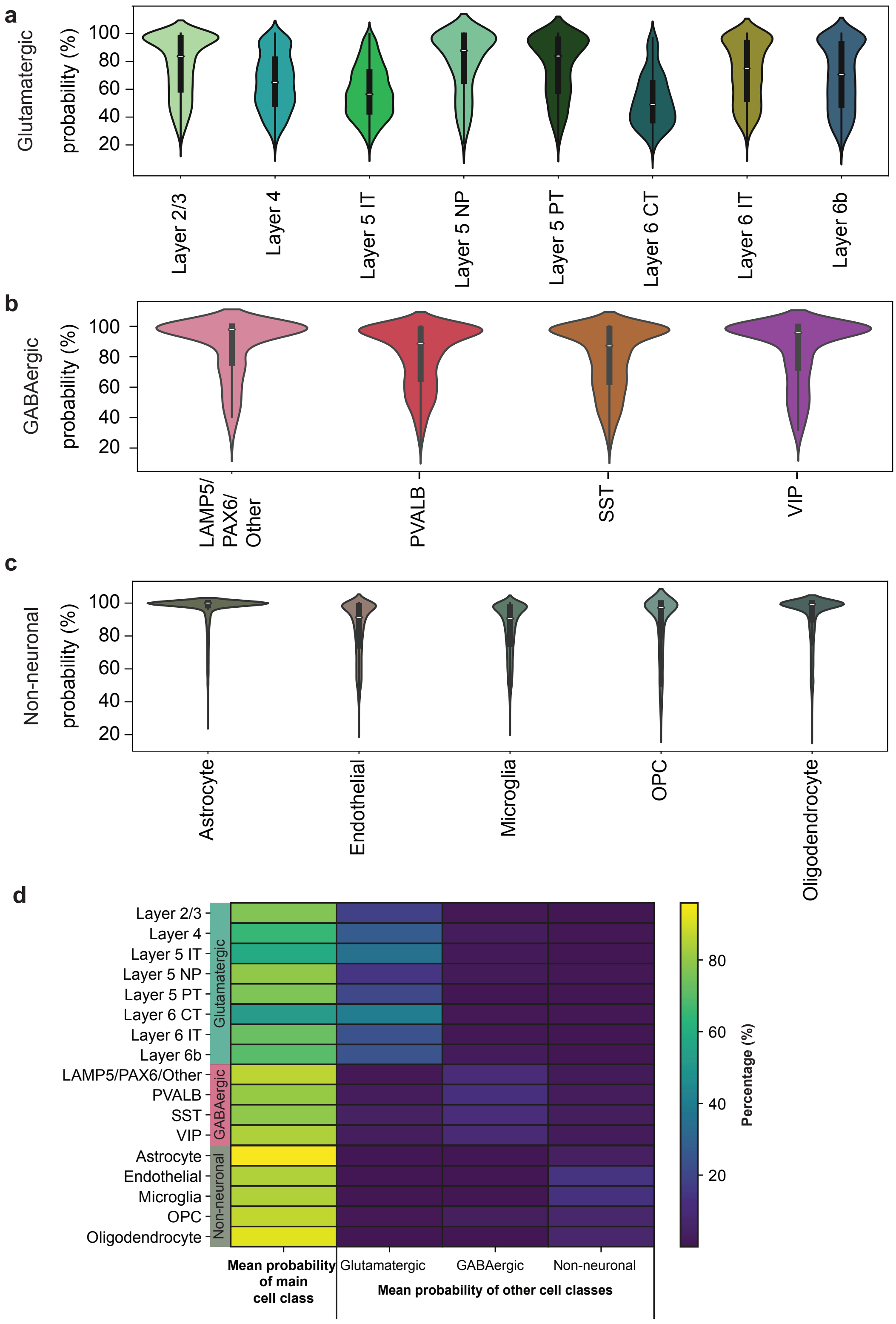


**Supplementary Figure 3 | Level of certainty for each cell subclass.**

(**a**) Violin plots of the highest probability in each pie chart from pciSeq for glutamatergic cells. The mean certainty for glutamatergic cells is 68.42 ± 0.23%.

(**b**) Same as (a) for GABAergic cells with a mean certainty of 81.42 ± 0.31%.

(**c**) Same as (a) for non-neuronal cells with a mean certainty of 89.38 ± 0.07%.

(**d**) Heatmap of the probability distribution in the pie charts for each cell subclass. Data is represented as mean value. The mean of the highest probability (largest pie) for each cell subclass is highlighted in the first column. The mean of the other probabilities (smaller pies) are shown in the remaining three columns and grouped for glutamatergic cells, GABAergic cells and non-neuronal cells. This highlights that cells mostly share pie charts with cell types of the same subclass, i.e. glutamatergic cells with other glutamatergic cells, GABAergic cells with other GABAergic cells and non-neuronal cells with other non-neuronal cells.

#### Supplementary Figure 4

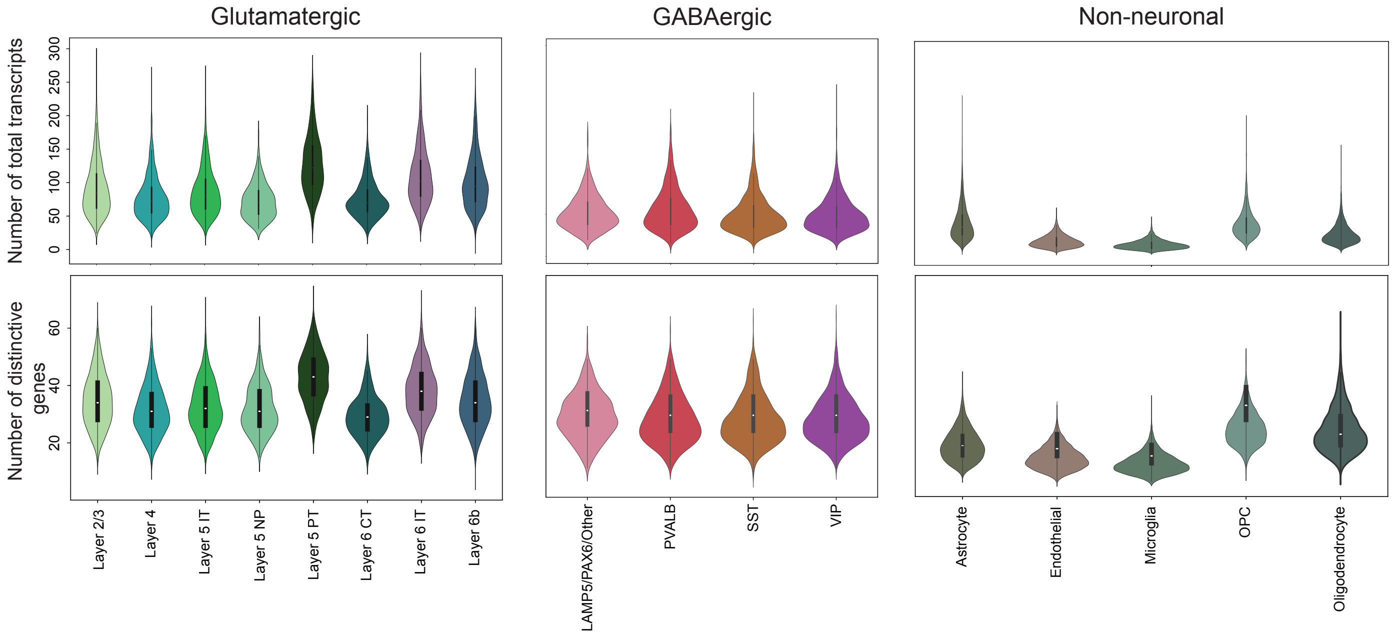


**Supplementary Figure 4 | Transcript and gene quantification in single cells.**

Top: Violin plots of the number of total transcripts in glutamatergic, GABAergic and non-neuronal cells. Bottom: The number of distinct genes in glutamatergic, GABAergic and non-neuronal cells.

### Supplementary Figure 5

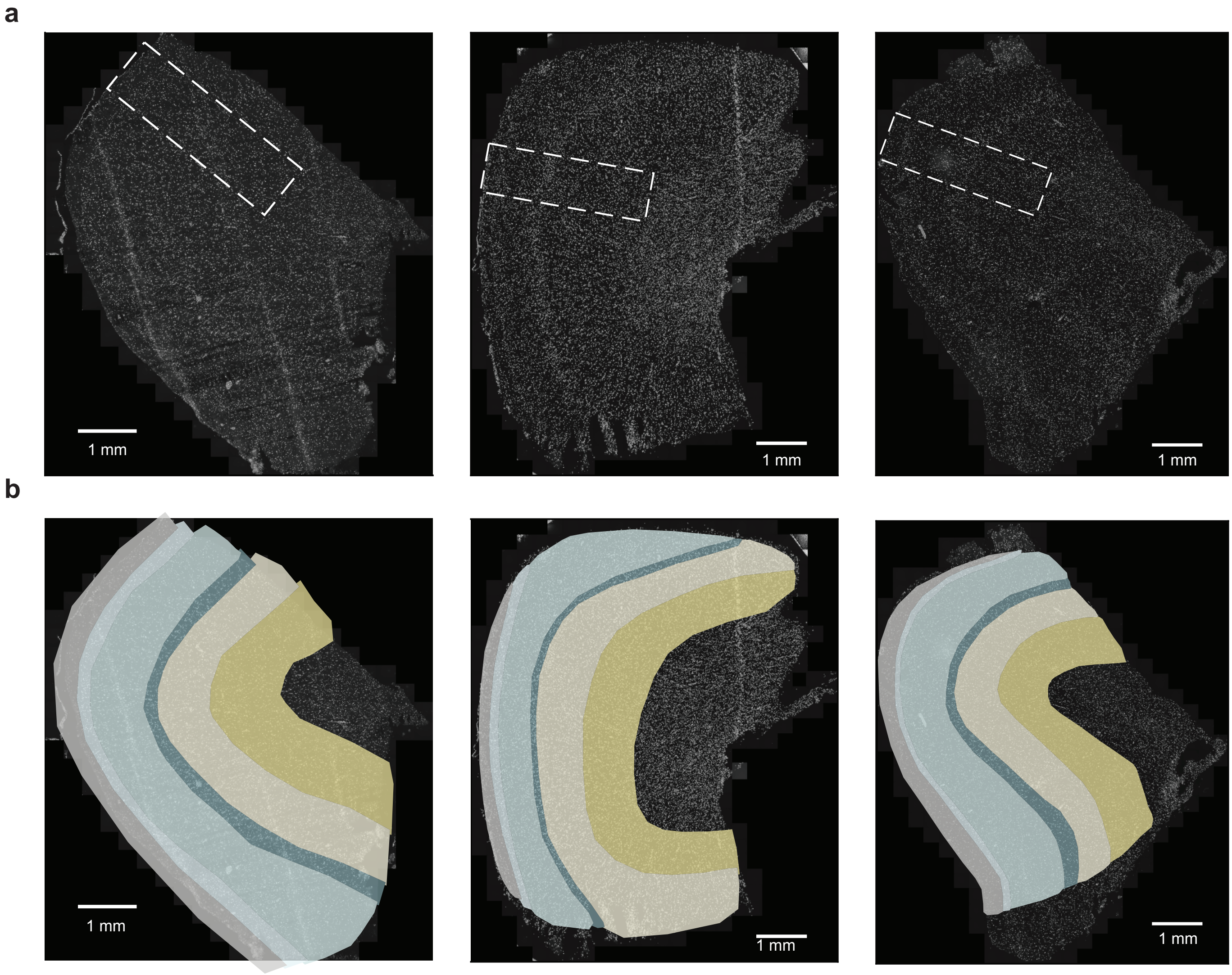


**Supplementary Figure 5 | Regions of interest and annotated neocortical layers in the human temporal lobe sections.**

(**a**) Region of interest on top of DAPI-stained sections for Sections A-C (left to right).

(**b**) Annotated neocortical layers on top of DAPI-stained sections for Sections A-C (left to right).

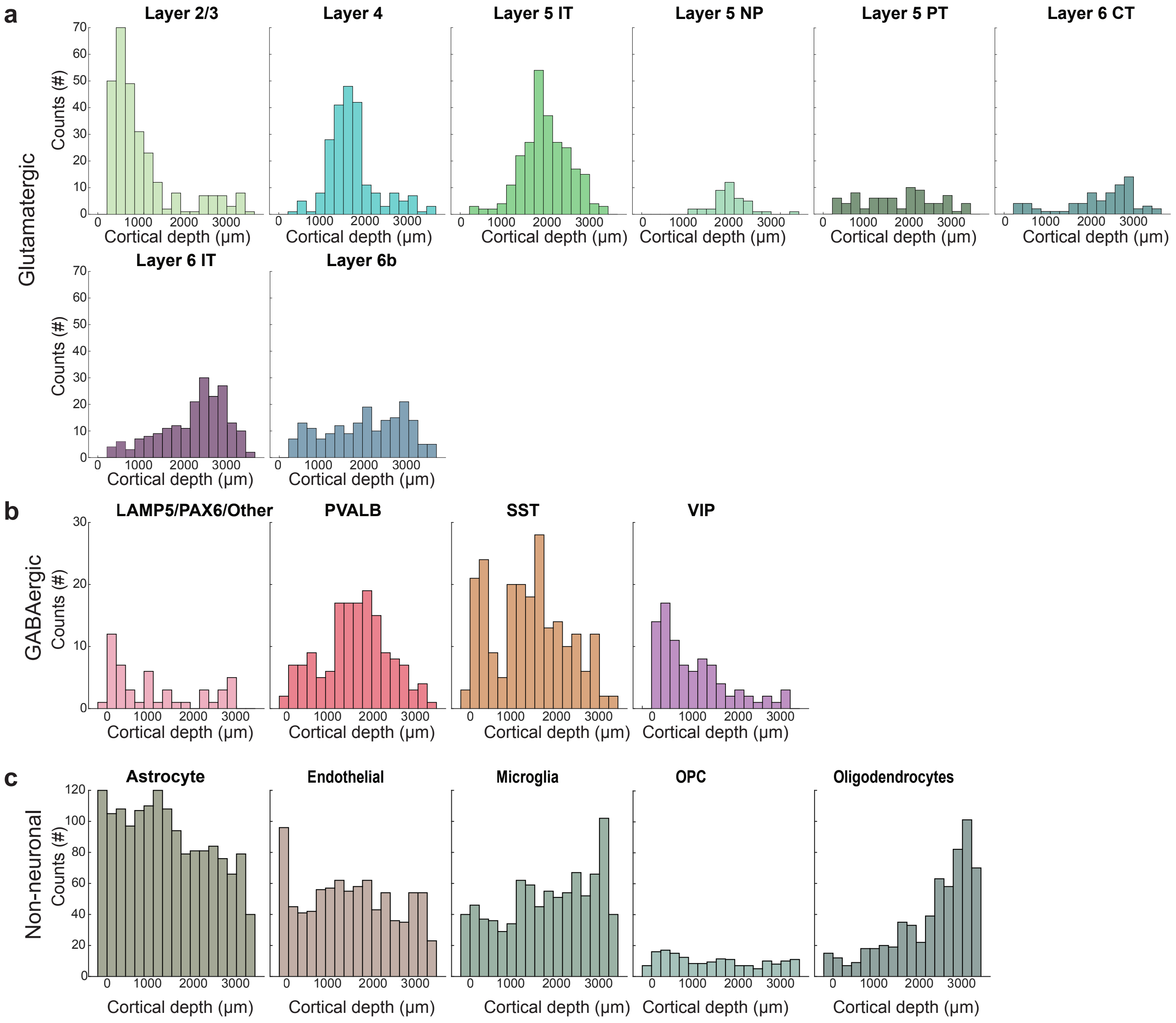


**Supplementary Figure 6 | Cortical profiles of glutamatergic, GABAergic and non-neuronal cells.**

(**a**) The cortical depth profile (1 bin = 200 µm) of the glutamatergic cell subclasses.

(**b**) Same as (a) for the GABAergic cell subclasses.

(**c**) Same as (a) for the non-neuronal cell subclasses.

#### Supplementary Figure 7

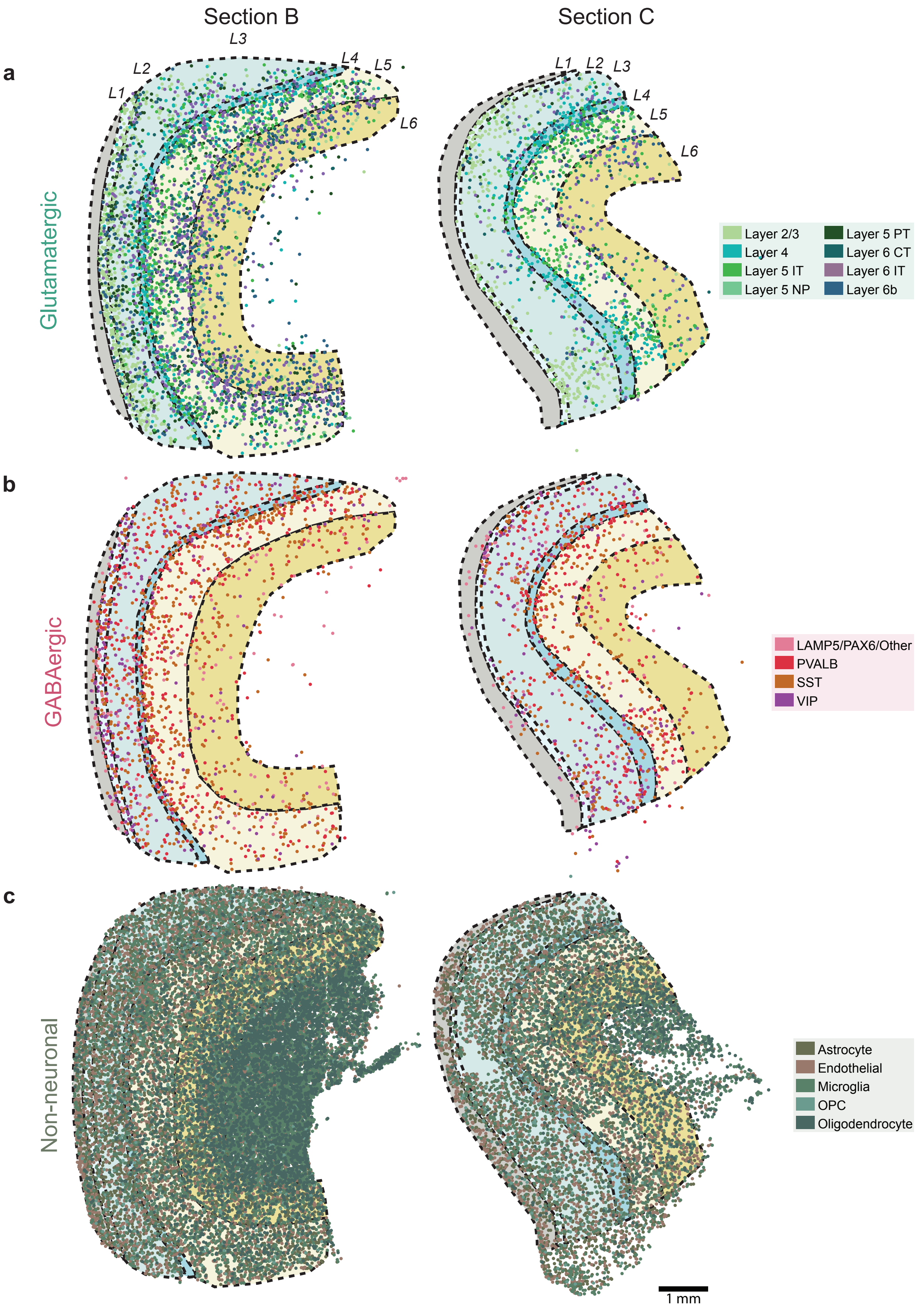


**Supplementary Figure 7 | Cell locations and annotated neocortical layers.**

(**a**) Location of glutamatergic cells colored by the most probable cell subclass (Section B (left) and C(right)). In the background, the annotated neocortical layers (L1-6) are plotted.

(**b**) Same as (a) for GABAergic cell subclasses.

(**c**) Same as (a) for non-neuronal cell subclasses.

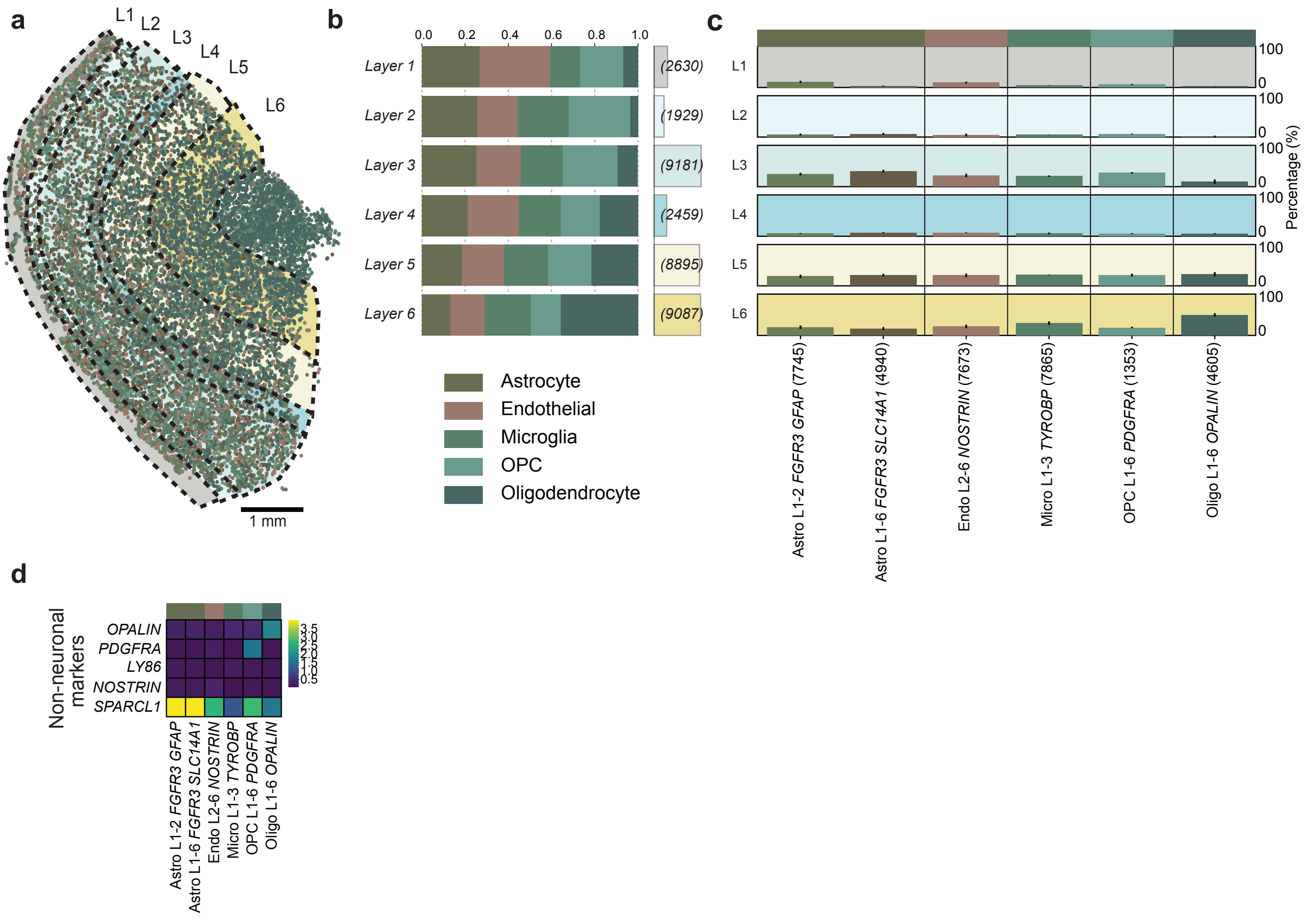


**Supplementary Figure 8 | Layer-specificity and molecular composition of non-neuronal cells.**

(**a**) Location of non-neuronal cells colored by the most probable cell subclass and annotated neocortical layers (L1-6).

(**b**) The within-layer relative distribution of non-neuronal cells and the number of cells counted for each layer in brackets.

(**c**) Across-layer distribution of non-neuronal cell types. The colored bars represent the relative proportion of each cell type in each layer (L1-6).

(**d**) Mean log2-transformed expression of known non-neuronal marker genes.

### Supplementary Figure 9

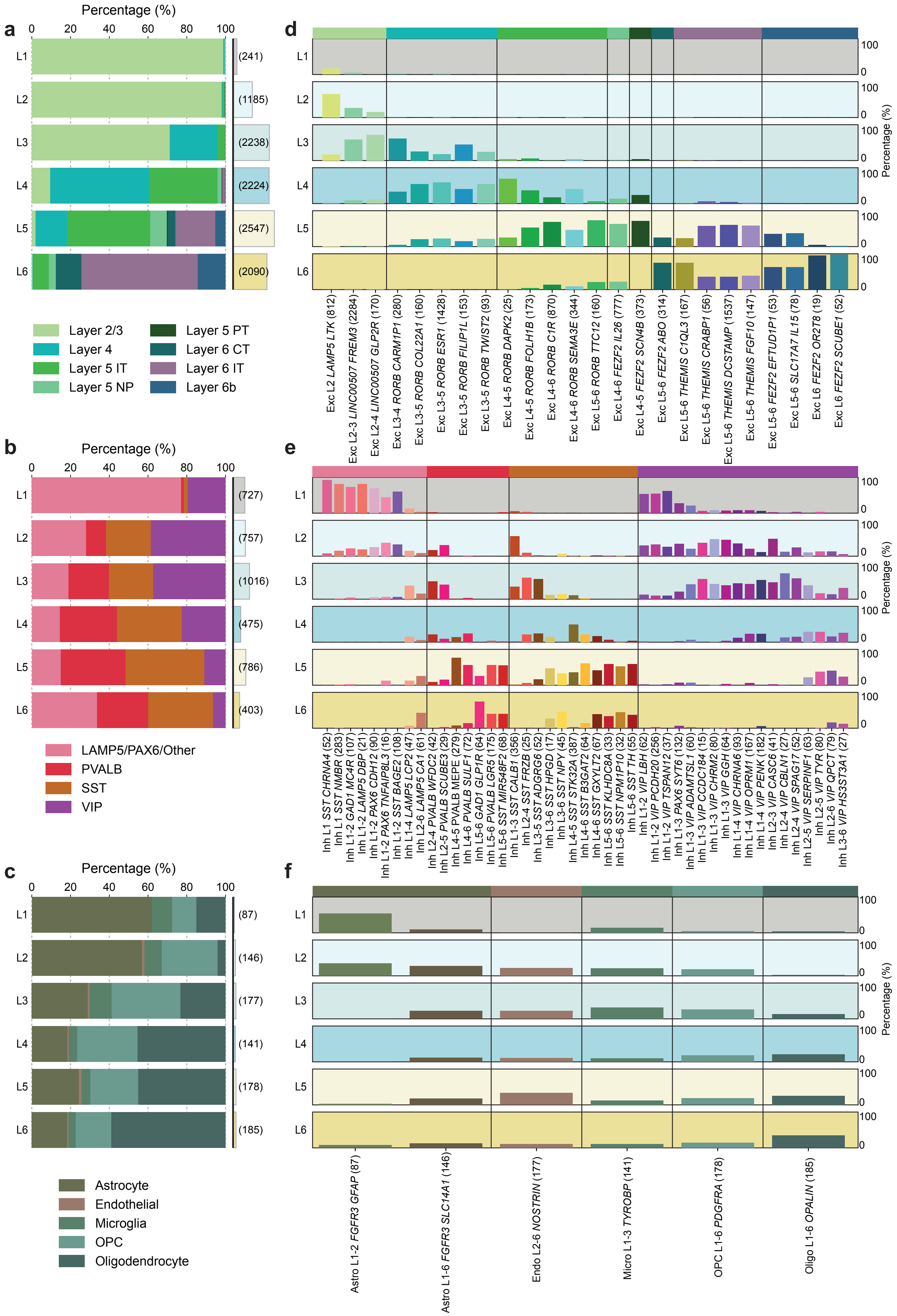

**Supplementary Figure 9 | Occurrences of glutamatergic, GABAergic and non-neuronal cells separated by layers in the human middle temporal gyrus from single-nucleus RNA-sequencing (snRNA-seq) data.**

(**a**) Single-nucleus RNA-sequencing (snRNA-seq) data from Hodge *et al.*[4] The within-layer relative distribution of glutamatergic cells and the number of cells counted for each layer in brackets.

(**b**) Same as (a) for GABAergic cells.

(**c**) Same as (a) for non-neuronal cells.

(**d**) Across-layer distribution of glutamatergic cell types. The colored bars represent the relative proportion of each cell type in each layer (L1-6).

*(***e**) Same as (d) for GABAergic cell types.

(**f**) Same as (d) for non-neuronal cell types.

### Supplementary Figure 10

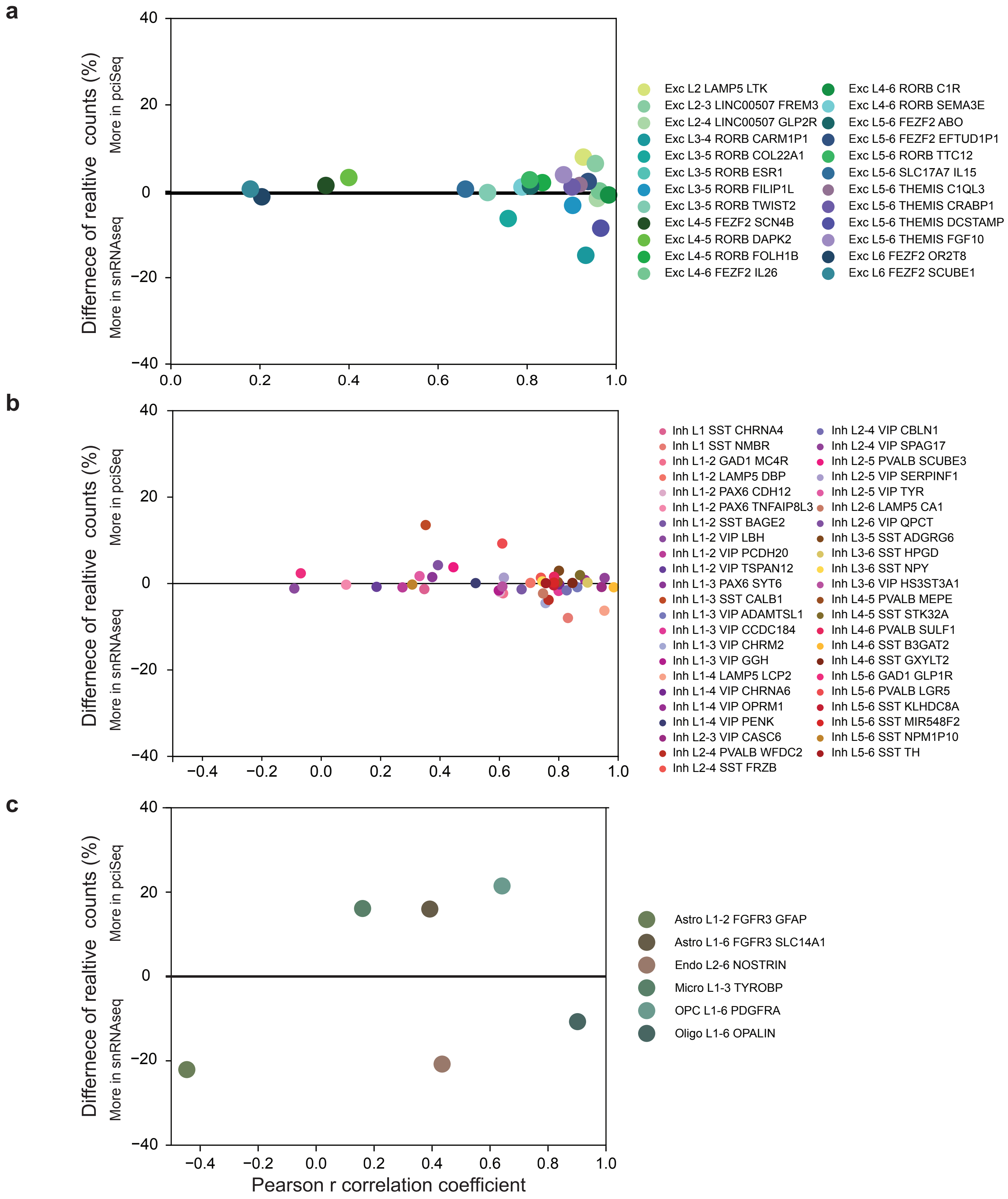


**Supplementary Figure 10 | Comparison of cell layer distribution and relative occurrences of cell types between pciSeq and snRNA-seq data.**

(**a**) Scatter plot of Pearson correlation coefficient on the X-axis and the difference in relative occurrence between pciSeq and snRNA-seq data on the Y-axis for the glutamatergic cell types.

(**b**) Same as (a) for GABAergic cell types.

(**c**) Same as (a) for non-neuronal cell types.

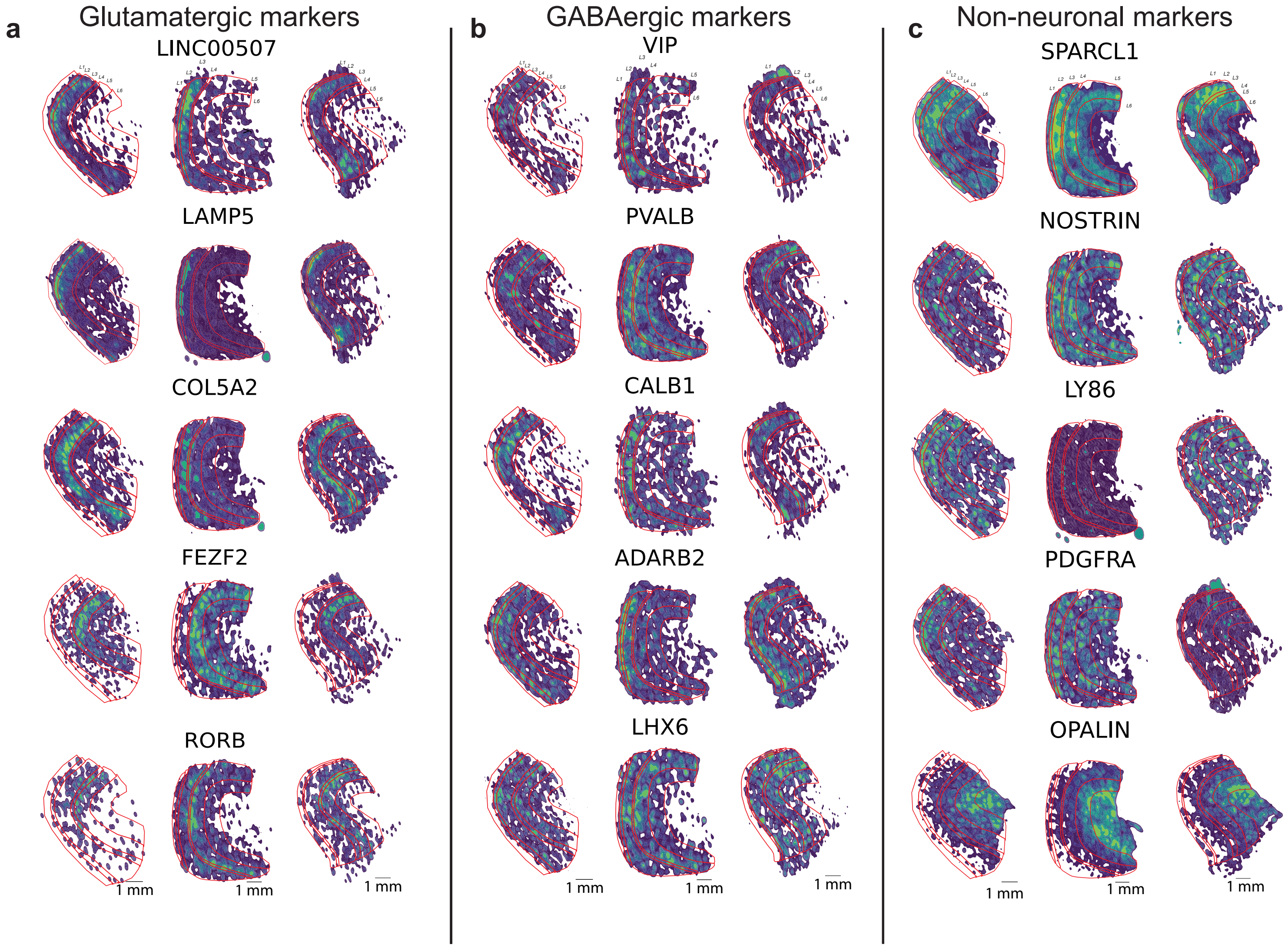


**Supplementary Figure 11 | Gene expression of known glutamatergic, GABAergic and non-neuronal marker genes.**

(**a**) Heatmap of the expression of known glutamatergic marker genes in the human temporal lobe sections. Layers are outlined in red to highlight the region specific expression.

(**b**) The same as (a) for GABAergic marker genes.

(**c**) The same as (a) for non-neuronal marker genes.
